# Electroencephalographic power spectrum and intersubject correlation on acoustic stimulation with modes of Indian music: a randomized controlled trial

**DOI:** 10.1101/2022.12.09.519709

**Authors:** Kirthana Kunikullaya U, Arun Sasidharan, Vijayadas, Radhika Kunnavil, Jaisri Goturu, Nandagudi Srinivasa Murthy

**Affiliations:** Univ Rennes, Inserm, EHESP, Irset (Institut de Recherche en Santé, Environnement et Travail), UMR_S 1085, F-35000 Rennes, France; The Maastricht Multimodal Molecular Imaging Institute, Maastricht University, 6229 ER Maastricht, The Netherlands; Center for Consciousness Studies (CCS), Department of Neurophysiology, National Institute of Mental Health & Neuro Sciences (NIMHANS), Off Hosur Road, Bangalore, Karnataka, India –560029; Department of Physiology, Ramaiah Medical College, MSR Nagar, MSRIT Post, Bangalore, Karnataka, India – 560054; National Institute of Unani Medicine, Under Govt. of India, Kottigepalya, Magadi Main Road, Bengaluru – 560091; Department of Physiology, International Medical School, MSR Nagar, MSRIT Post, Bengaluru 560054, Karnataka, India; Department of research and patents, Gokula Education Foundation, MSR Nagar, MSRIT Post, Bangalore, Karnataka, India – 560054

**Keywords:** Indian music, melodic modes, electroencephalogram, inter-subject correlation, correlated component analysis

## Abstract

**Background:** Music not just entertains an individual but causes changes in the frequency spectrum of the brain waves and cognition that are recognizable using signals obtained through electroencephalography (EEG). EEG studies on the effect of passive listening to music have predominantly used multi-instrumental western classical music as an acoustic stimulus with very few analyzing solo instrumental Indian music, and thus in the current study Indian modes (*Hindustani ragas*) were used. The study aimed to investigate overall power spectral changes on EEG and specifically, those changes that show high inter-subject correlation (ISC) on passive listening to three different Indian modes as acoustic intervention, in comparison to control stimuli, heard for 10 minutes.

**Material & Methods:** A randomized control triple-blind trial with 4 groups (three music intervention groups and a control group; n=35 each) was conducted while undergoing EEG recording. The music intervention groups listened to 10-minute audio of one of the three different modes (namely *raga Miyan ki Todi, raga Malkauns*, and *raga Puriya*), while the control group received predominant silence with few natural sounds interspersed. EEG data before, during, and after acoustic interventions were first evaluated for electrode-level power changes in standard spectral bands (delta, theta, alpha, beta1, beta2, and gamma). To understand spectral power changes more specific to music listening, a novel component-level analysis was also done, where the raw spectral data were grouped into the three most prominent components (C1, C2 & C3) based on spatiospectral consistency across subjects (correlated component analysis or CorrCA) and their ISC scores were also computed. For statistical analysis, we applied a hierarchical general linear model with cluster statistics to the electrode-level data and robust ANOVA with post hoc tests to the component-level data.

**Results:** In electrode level analysis, the group listening to *raga Malkauns* showed a significant increase in gamma power in the left frontal regions during the intervention. While the group listening to *raga Puriya* showed a right frontoparietal decrease in delta power, *raga Miyan ki Todi* showed a frontal increase in beta1 power after the intervention. In component-level analysis, C1 was globally distributed low-frequency activity, C2 was posteriorly dominant alpha-beta1 activity, and C3 was peripherally dominant broad-band activity, consistent between groups. Besides agreement with electrode-level findings, the most prominent component-level finding was a decrease in C1 power and an increase in C2 power shown by *raga Malkauns* (strong both during and after intervention) and *raga Miyan ki Todi* (strong during and weak after intervention), whereas *raga Puriya* showed only a weak decrease in C1 (after intervention), compared to control group. ISC scores were comparable between groups, except for *raga Puriya*, which showed a marginal drop for C3 after the intervention.

**Conclusions:** Reduction in globally distributed low-frequency activity and increase in posterior dominant alpha-beta1 activity may be characteristic of passive listening to relaxing Indian modes, which may persist even after the listening period. Among the modes, *raga Malkauns* showed this effect most prominently, followed by *raga Miyan ki Todi* and least by *raga Puriya*. As the increase in posterior alpha and low beta power is associated with default mode network (DMN) activity and a decrease in delta power with positive emotional memory, the spectral pattern we observed may indicate observing positive autobiographical memory while listening to musical scales and thereby contributing to a relaxing experience. Further studies that also include phenomenological reports are highly recommended to be taken up to support these findings, and thus build a scientific foundation for the use of Indian music in medicine.

**Graphical Abstract:** 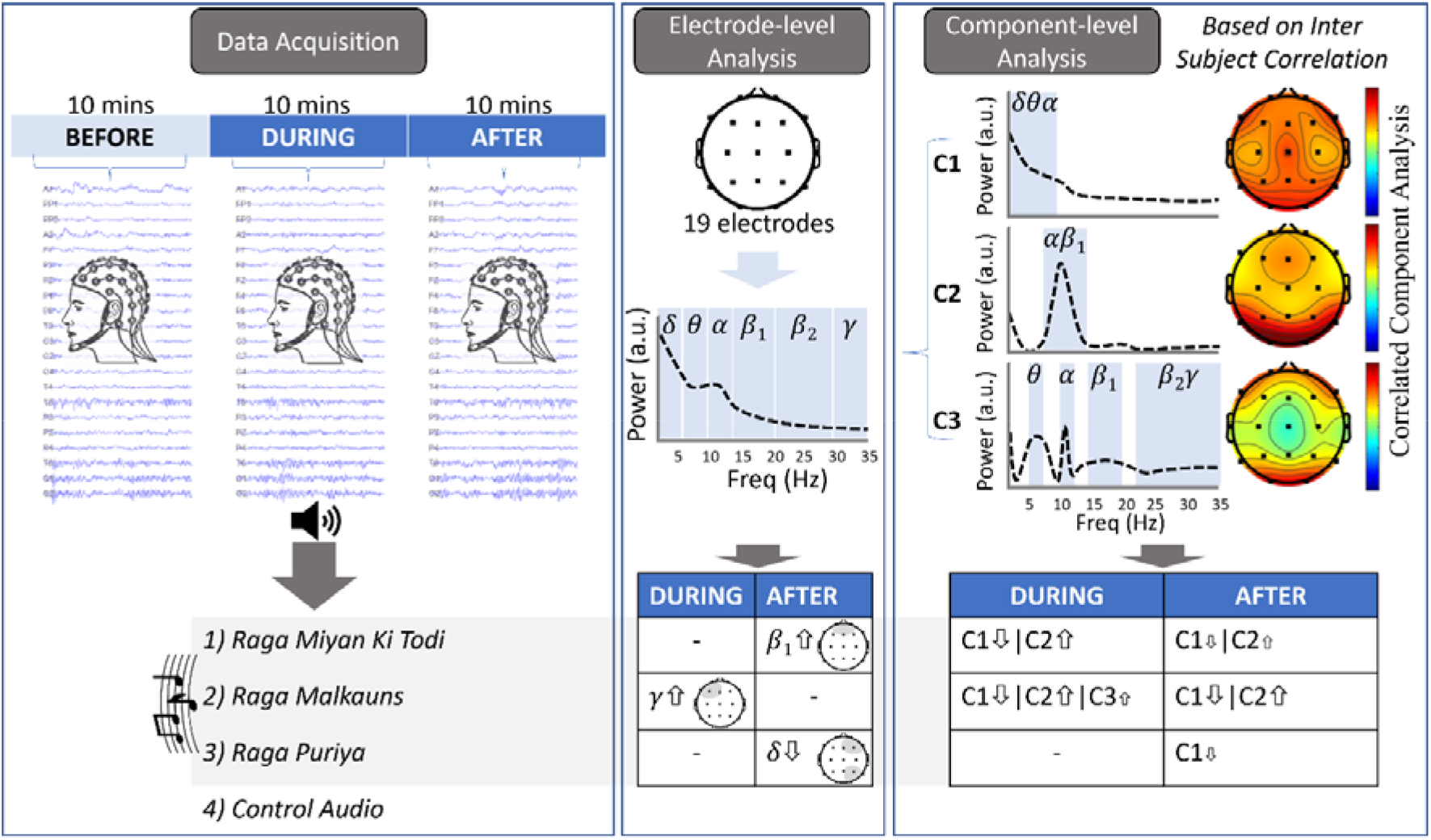

**Impact Statement:** In this manuscript on EEG power spectral changes, correlated component analysis, and Intersubject correlation, we found a reduction in globally distributed low-frequency activity and an increase in posterior dominant alpha-beta1 activity during listening to Indian modes of music, in comparison to the control group. Mode-specific changes in correlated components, indicating default mode network activity and probable activation of autobiographical memories were observed.

## Introduction

Health benefits of exposure to natural environments include reduction of stress and the creation of a positive attitude towards resource conservation, through stimulation of visual, auditory, or multisensory systems (1–3). One of the universal ways a human connects with the environment is through engagement, which is defined behaviorally as a commitment to attend to the stimulus. Similarly to other sensory experiences, acoustic stimulus, especially music, engages a person and helps generate emotions (4,5). Music not just entertains an individual but affects cognition (6,7), including emotion modulation (8,9) and stress reduction (10–12). These responses can be recorded using subjective [emotion questionnaires, valence, and arousal (13) recordings] and objective [electrocardiography (14), EEG, fMRI measures (15,16)]. In the current study, we used EEG to evaluate the effect of the acoustic stimulus with music.

Listening to Indonesian Gamelan music increased EEG beta power, bilateral temporal region blood flow, and freshly recruited the posterior portions of precuneus bilaterally (cerebral blood flow positively correlated with the beta power in these regions) (17), indicating music-evoked memory recall or visual imagery (18). Listening to instrumental music (*raga Desi Todi*), for 20 days (30 min each day) reduced anxiety and increased alpha power on EEG (19). Using multifractal detrended fluctuation analysis (DFA) of EEG, it was shown that acoustic stimulus with the drone instrument (tanpura) increased the complexity of the frontal alpha and theta power (20). Music therapy for stress reduction increased posterior theta power and decreased mid-frontal beta and posterior alpha power, along with a reduction in the degree of anxiety (21). It is important to note that several studies have studied the acute effects of music and have used durations ranging from a few seconds to about 3 minutes. Longer durations of exposure to music have been explored less frequently in research, except in follow-up studies, active music interventions, or when used for therapeutic purposes. In our previous study, we observed an overall effect on the EEG power spectrum and on listening to music (for ten minutes). Furthermore, we noted significant temporal differences within the time intervals (every 2 minutes) that were analyzed as the music was heard over this duration (16). Therefore, music being a time-based stimulus that unfolds over time, it is important to analyze these brain wave changes with time as an important factor.

Further, when a set of individuals are exposed to the same sensory stimulus, few individuals will have typical experiences and they are said to be engaged with the stimulus. However, the experience of few remains atypical, which is said to be due to the typicality of their stimulus-evoked brain activity (22). This can be measured by inter-subject correlation (ISC), which evaluates the similarity of an individual’s brain over some time with that of another individual or a group, in a given region of the brain. A detailed tutorial on ISC is provided in (23). Predominantly ISC is used with fMRI or EEG data from individuals visualizing a moving stimulus (movie clipping) (22,24–28). As music is a time-based stimulus, along with other factors, it is important to understand engagement and ISC while music is used as an intervention. ISC of neural responses is said to be well-suited for measuring musical engagement (29). A recent study found that ISC reduced after repeated listening to familiar music compared to unfamiliar classical music pieces, lasting for about 60 – 90 seconds (30).

Although previous studies have investigated brain responses during continuous listening to relatively simple musical stimuli or controlled auditory paradigms (31–33), there is a dearth of parallel studies that examine how the human brain processes the multitude of musical features during passive listening to naturalistic musical stimuli. Until now, there has been little discussion on changes in EEG power and ISCs on receptive listening to different melodic modes over a longer duration. Therefore, in this investigation, the main objective was to analyze the spectral variations of EEG power and the cortical entrainment to music during receptive listening to three different melodic modes, in addition to ISCs. For this study, solo instrumental Indian music modes were used, without percussion or lyrics, ensuring uniformity of the intervention. As repetition is said to reduce ISC values (25–27), in the current study, acoustic stimuli were not repeated within the same individual or between groups. We hypothesized that due to the repetition of the phrases during the elaboration of a given melodic mode (in the given duration) the overall ISC may reduce.

## Materials and Methods

### Study design & ethics

This was a prospective randomized controlled trial with a triple-blind design (part of a larger trial (34)) with 140 participants who were randomly divided into 4 groups, with 35 participants in each group. Each group received one of the acoustic interventions, group A received *raga Miyan ki Todi*, group B received *raga Malkauns* and group C received *raga Puriya*, with group D as a control group. We recorded EEGs during three tasks: silence, music listening, and silence (each lasting 10 minutes) and compared them with the control (no music). The study period ranged from 2019 to 2021 (June 2019 - first recruitment and February 2021 - last recruitment). The data presented here were taken from a larger experiment (full trial protocol: NCT03790462 on clinicaltrials.gov.in). The research was conducted following the Declaration of Helsinki guidelines. The study protocol was approved by the institutional scientific committee on human research and the ethical review board (Reference: MSRMC/EC/2017, dated: 25/07/2017).

### The basis for sample size

The sample size was calculated based on a previous study where the change in State Trait Anxiety Inventory-6 (STAI-6) anxiety median and IQR scores was from 33.3 (23.3– 41.7) before music intervention and to 30 (20–40) after music intervention. Considering the minimum difference of 4 units in the STAI score, with an effect size of 0.7, power of 85%, and an alpha error of 5%, the sample size was calculated to be 35 in each group (35).

### Recruitment

The recruitment of participants, the randomization technique, and the process of EEG recordings are explained in detail in (16,34). Briefly, healthy right-handed participants aged 18 to 30 years, of any sex, fluent in English, with normal hearing, without cognitive or decisional impairments, and who were not smokers or alcoholics were invited to participate in the study, and those who responded were asked to complete an online questionnaire. Participants with any past or current medical or surgical disorders, hearing impairment, and self-reported BMI of over 30 kg/m^2^ were excluded from the study [Figure 1]. The rest of the participants were invited to the lab for further recordings.

**Figure 1:**
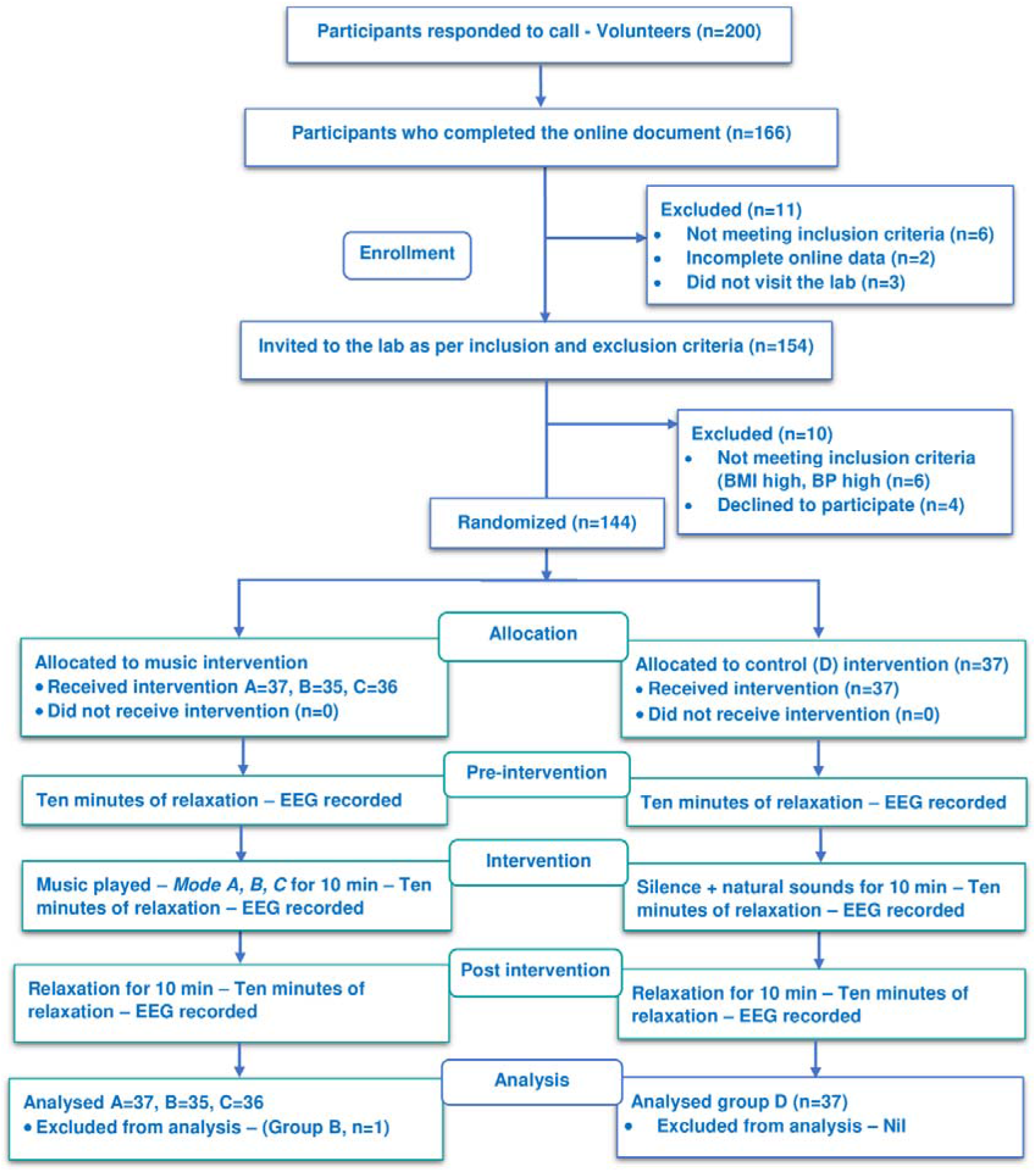
Consort diagram of participant recruitment, distribution, and follow-up Intervention

### Randomization

A simple randomization technique was used to randomly select participants into four groups (Fig 1). The random numbers were generated using MS Excel (4 sets of 35 each) and sealed in an opaque envelope. The serial number of the participants was written at the top of the envelope. The envelope was opened by the research assistant after the baseline assessment of each participant and then they were assigned to their respective arms. The participants knew understood that they were in the music intervention group once the intervention began, though they did not know the mode. All the investigators who did the outcome assessments were blinded to the interventions.

Three melodic scales (modes/*ragas*) were chosen and implemented as interventions. Indian classical melodic scales/modes (*ragas*) *Malkauns* (pentatonic: C, E⍰, F, A⍰, B⍰), *Puriya* (hexatonic: C, D⍰, E⍰, F#, A, B) *& Miyan ki Todi* (heptatonic: C, D⍰, E⍰, F#, G, A⍰, B) were chosen based on our previous work [Table 1], where we standardized the music (10,11,34).

**Table 1:**

| Svara / Note | Hindustani name | Staff note | Western scale Interval name |
| --- | --- | --- | --- |
| **Raga Miyan ki Todi (Scale A) (heptatonic, G appears in descent)** |  |  |  |
| S | Shadja | C | Perfect unison |
| r | Komal Rishab | D$\flat$ | Minor second |
| g | Komal Gandhar | E$\flat$ | Minor third |
| M | Tivra Madhyam | F$\sharp$ | Augmented fourth |
| P | Pancham | G | Perfect fifth |
| d | Komal Dhaivat | A$\flat$ | Minor sixth |
| N | Shuddha Nishad | B | Major seventh |
| **Raga Malkauns (Scale B) Ascent and descent same – pentatonic** |  |  |  |
| S | Shadja | C | Perfect unison |
| g | Komal Gandhar | E$\flat$ | Minor third |
| m | Shuddha Madhyam | F | Perfect fourth |
| d | Komal Dhaivat | A$\flat$ | Minor sixth |
| n | Komal Nishad | B$\flat$ | Minor seventh |
| **Raga Puriya (Scale C) C, D$\flat$, E, G$\flat$, G, A/A$\flat$, B (hexatonic)** |  |  |  |
| S | Shadja | C | Perfect unison |
| r | Komal Rishab | D$\flat$ | Minor second |
| G | Shuddha Gandhar | E | Major third |
| M | Tivra Madhyam | F$\sharp$ | Augmented fourth |
| D | Shuddha Dhaivat | A | Major sixth |
| N | Shuddha Nishad | B | Major seventh |

The music was tuned to a frequency of 329.63 Hz (the tonic or ‘*Sa*’ at pitch E). The music used for this study was 10 to 11-minute-long instrumental (Flute/*Bansuri*) music recorded by an eminent musician (exclusively for the present study), playing the improvisation in the respective scales [named *alaap* (11,34,36) in Indian music]. We chose the flute as the instrument to play the melodic scales, as only musical components such as pitch, intensity, rhythm, and timbre would be present, without the lyrical and percussion components (37–45). Participants in group D (the control group) did not have any intervention except a few natural sounds that were played at the same volume levels once every 2 minutes for 10 seconds, similar to our previous study (16). The acoustic stimulus was coded as A, B, C & D by a person uninvolved in the project. We instructed the participants to listen to this with their eyes closed, and minds relaxed, for the duration it was played (10 to 12 minutes). All participants listened to the acoustic intervention through headphones [considered ideal according to review (46)], connected to a laptop, at a uniform volume (50%).

### Process of recordings

Recordings of EEG were done in a supine position with eyes closed, with the first five minutes utilized for attaching EEG electrodes. The participant was asked to relax with eyes closed for the next 30 – 35 minutes when the EEG was continuously recorded, with the event marking artifacts such as eye movement, jaw movement, and acoustic intervention played through headphones, in the mid 10 minutes (Fig 2). We concentrated on the effects that would naturally occur during acoustic stimulation, without the participation of the participants in any particular cognitive task (47). Subsequently, the participants heads were cleaned and relieved. The recordings were made using a 19-channel EEG system (Galileo Mizar Lite, EB Neuro, Italy), with silver chloride electrodes (Ag-Cl) placed on the scalp after the 10-20 international system of electrode placement [active electrodes were placed in Fp1, Fp2, F7, F3, Fz, F4, F8, T3, C3, Cz, C4, T4, T5, P3, Pz, P4, T6, O1, O2 and the reference electrode in the ear lobes (A1 & A2)].

**Figure 2:**
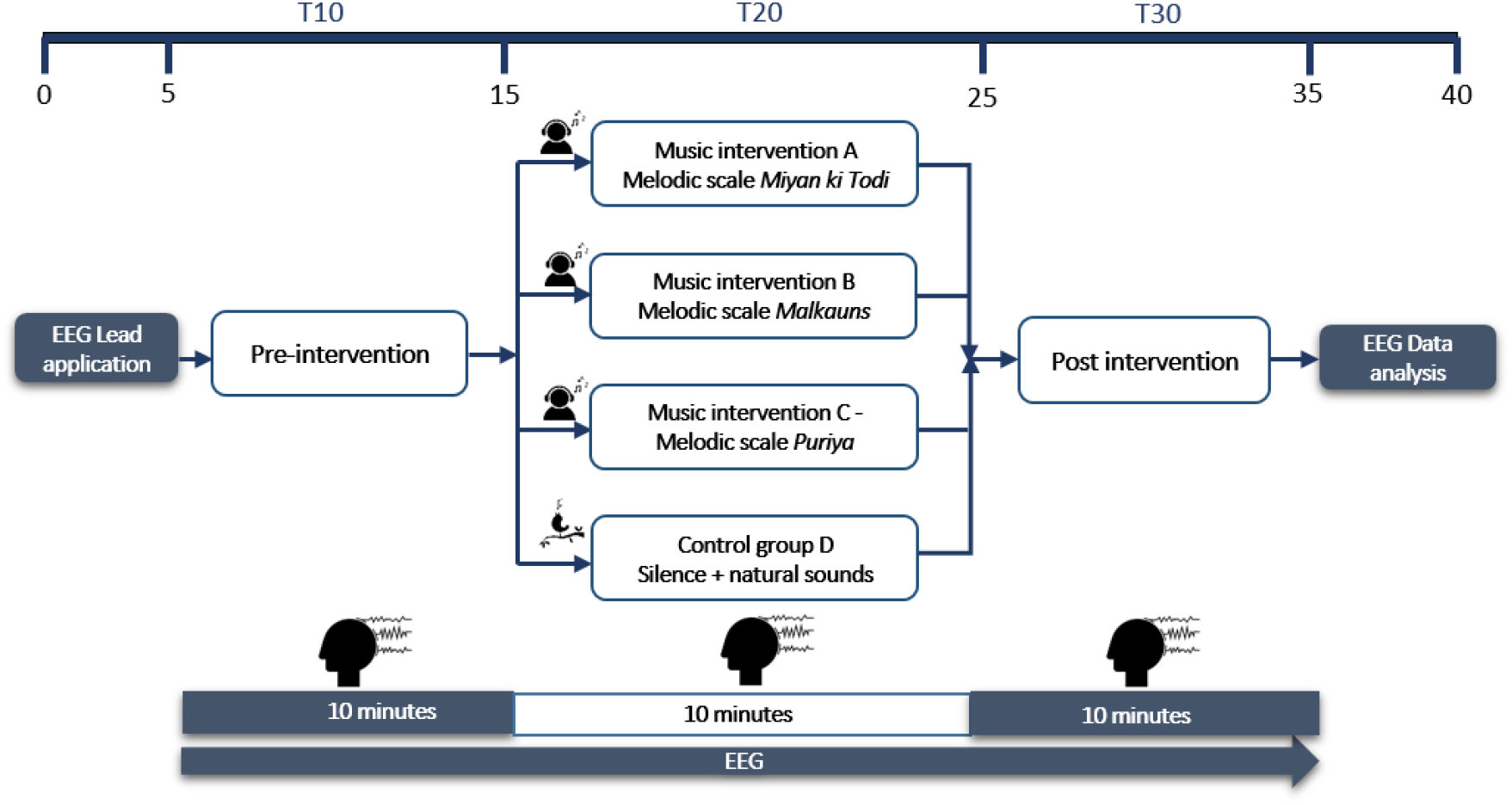
Flow chart describing the process of recording

### Electroencephalographic Data Analysis

Raw recordings of 3 conditions were marked before, during, and after interventions and stored in GalNT software (EB Neuro, Italy). These were then converted to the standard European Data Format (EDF). As the EEG data exported from the device did not have markers, a researcher manually recorded the timings for each of the 3 study conditions (i.e., Before, during, and after the intervention) in the device software. These timings were carefully corrected for data clipped off during acquisition pauses (there was at least one pause per subject). Using custom code written in MATLAB software, event marker files (.evt files required for the next step) were generated for each EEG file, denoting the 7-minute continuous segments within each of the 3 study conditions (skipped the initial 1 minute of each condition to avoid artifacts during transitions) (16). These data were then subjected to pre-processing using EEGLAB (48) functions (version: 2021). This included: 0.5-40Hz bandpass filtering, automated bad channel and bad segment removal by artifact subspace reconstruction (ASR) approach (49), bad channel spline interpolation, and average re-referencing - all done with custom written codes in MATLAB software (version: 2021b). The cleaned EEG data would be discontinuous with multiple short bad segments removed (varied across participants) and ‘boundary markers’ added to handle boundary effects during further analysis (which do not require long continuous EEG segments). Therefore, these were further visually inspected and only data from participants who had a total of at least 3 min of good quality EEG for all three portions were chosen for further analysis.

Band power was evaluated using FFT-based power spectral analysis on each of the EEG segments (with 2 seconds nonoverlapping Hanning windowed subepochs giving 0.5Hz resolution) and grouped in standard frequency bands (Delta: 1-4Hz, Theta: 4-7Hz, Alpha: 8-13Hz, Beta: 13-30Hz, and Gamma: 30-45Hz), for all 19 EEG channels, across all files. The gamma power was limited to <45Hz) due to the low sampling rate (128Hz) and potential filter effects in this frequency band.

As multi-channel EEG captures spatially distributed activity of prominent brain networks, instead of analyzing individual electrodes, we decided to use a linear combination electrode-level spectral activity that captures EEG spectral activity most correlated between participants. Correlated component analysis (CorrCA) helped achieve this and is conceptually based on canonical correlation analysis. This approach has been used in prior studies on time-domain data to extract multi-electrode EEG components related to music listening and video viewing (25,30). As our EEG segments were not well timed for intervention stimuli, we used frequency domain data (average power spectral data within each 10 min condition) for CorrCA. CorrCA also gives the forward model topography and time series of the components which could be used for spectral and other analyses that are typically done on single electrode data. For CorrCA, we used the code available at http://www.parralab.org/isc/. For the current analysis, we performed CorrCA on compiled spectral data from each group separately to capture the components most prominent to each intervention and selected the first three components per group with the highest ISC values at the group level. The subject-level total power across the spectra of each component was examined for the intervention effect between groups. Furthermore, the ISC values at the subject level were compared as a measure of engagement within the conditions. For statistical analysis, the data during and after the intervention were subtracted from the data before the intervention and then used for comparisons between groups. The results were also compiled into a Master Chart for further processing in a statistical tool using other physiological, psychological, and sociodemographic variables.

### Statistical analysis

Data were analyzed using SPSS software version 18.0 (SPSS Inc. Released in 2009. PASW Statistics for Windows, Version 18.0. Chicago: SPSS Inc.). The continuous variables were analyzed using descriptive statistics and the qualitative/categorical variables were analyzed using frequency and percentage. For statistical analysis of electrode-level EEG spectral data, we applied a hierarchical general linear model with cluster statistics to the electrode-level data using functions of the LIMO toolbox in MATLAB software. For statistical analysis of component-level spectral data and ISC scores, we applied robust one-way repeated measures ANOVA on trimmed mean followed by post-hoc Yuen’s trimmed mean test (20% trimming) (50) and p-values adjusted using Holm’s correction (51), using Jamovi software written in R language. Two-tailed p value ≤0.05 was considered statistically significant at a 5% level of significance.

## Results

General sociodemographic characteristics and other questionnaire data can be found in (34). Breifly all the sociodemographic characteristics were comparable between the groups, except educational status, which was adjusted for during physiological parameters analysis. There were no differences in familiarity to music or training in between the groups.

### Electrode-Level Band Power Changes Relative to Baseline

We calculated the power changes across the scalp for each subject, relative to the baseline power before the intervention, for each frequency band. These power changes were obtained as first-level beta values of a hierarchical general linear model approach described in the Methods section. During the intervention, the most prominent changes were a global decrease in alpha power for all intervention groups and a frontocentral increase in beta/gamma for two of the music intervention groups (*raga Malkauns* and *raga Miyan ki Todi)* (Fig 3). After the intervention, the most prominent change was a frontal decrease in delta power and a frontal increase in beta1 power in most groups (Fig 4).

**Figure 3:**
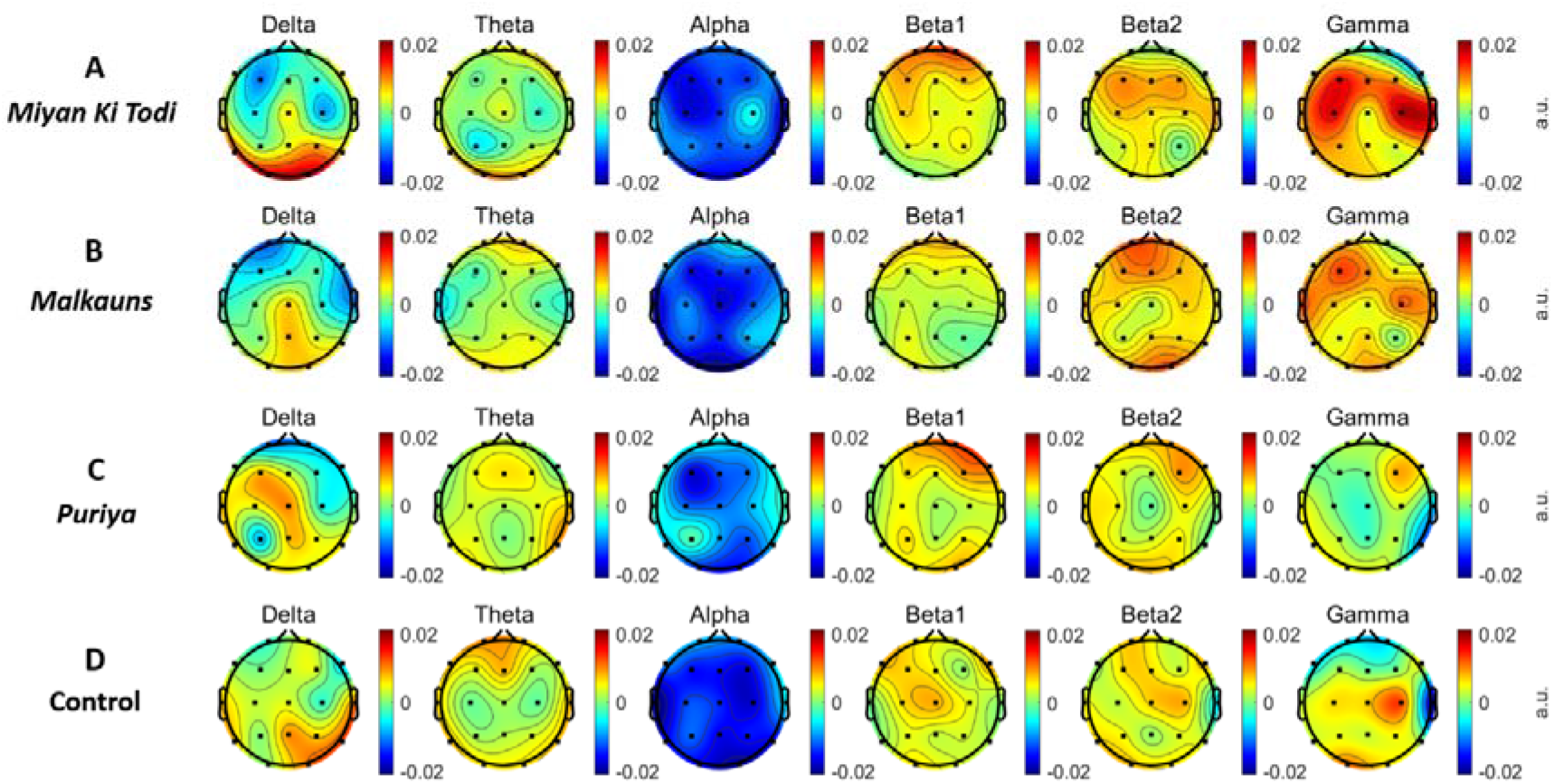
Scalp maps representing the average changes in electrode-level band power during intervention

**Figure 4:**
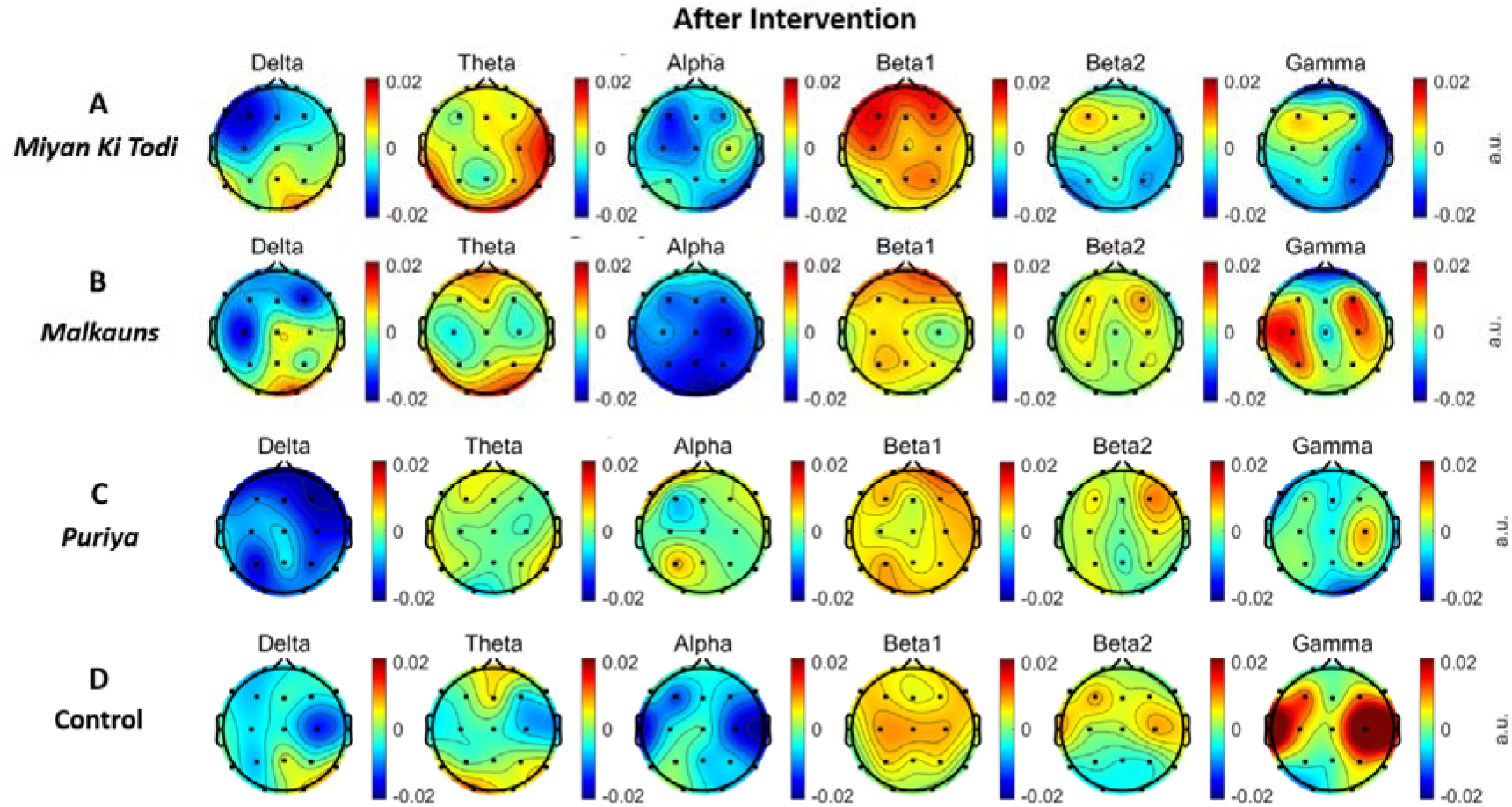
Scalp maps representing the average electrode-level band power changes after intervention

To evaluate the group differences, a second-level analysis was done where each music intervention group was statistically compared with the control group and cluster statistics (tfce) was employed to determine the significant changes. Based on this analysis, the group listening to *raga Malkauns* showed a significant increase in gamma power in the left frontal regions during the intervention (Fig 5). While the group listening to *raga Puriya* showed a right frontoparietal decrease in delta power and those to *raga Miyan ki Todi* showed a frontal increase in beta1 power, after the intervention (Fig 6).

**Figure 5:**
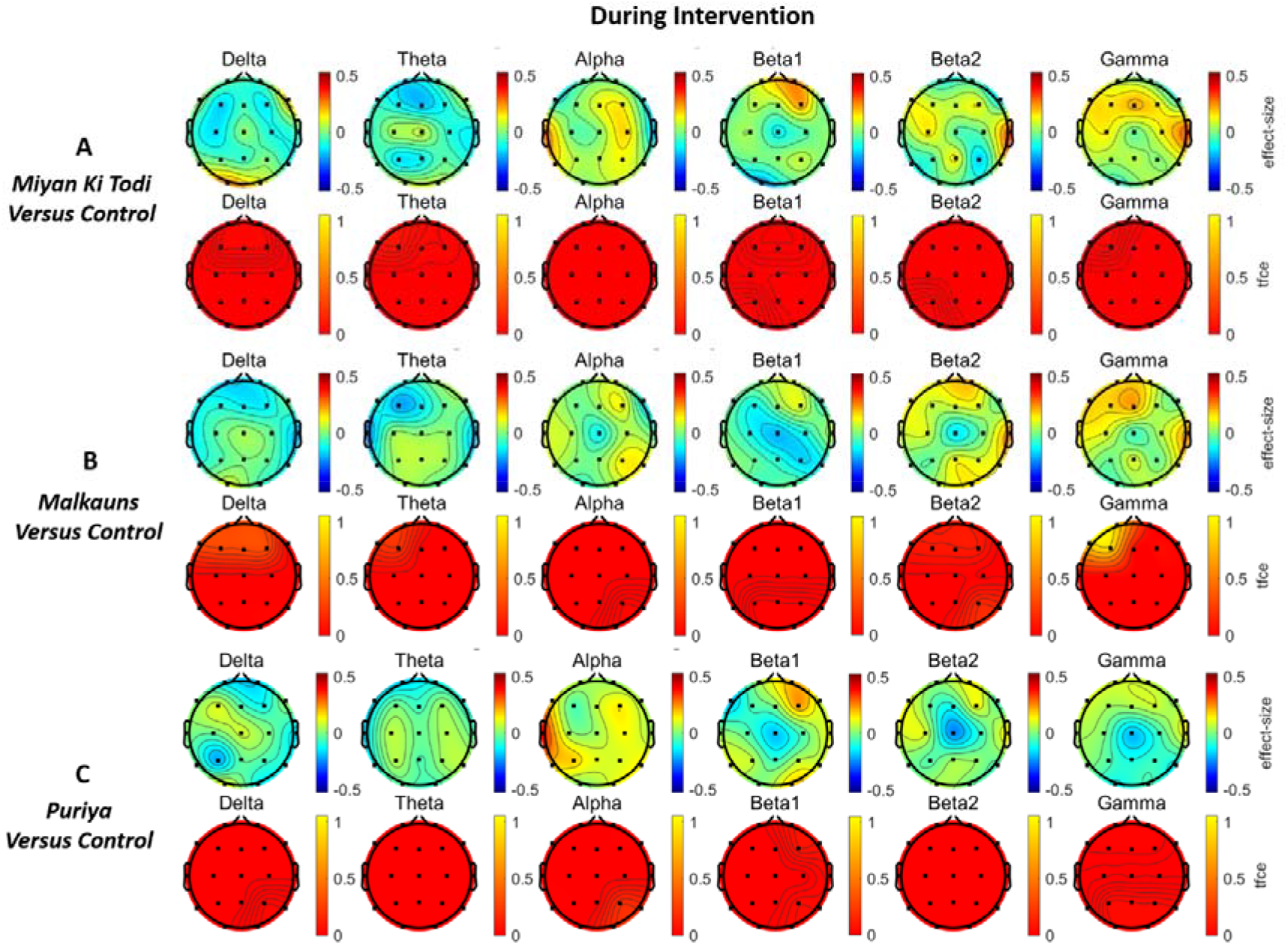
Scalp maps showing the differences between the groups in electrode-level band power between the intervention and control groups during the intervention.

**Figure 6:**
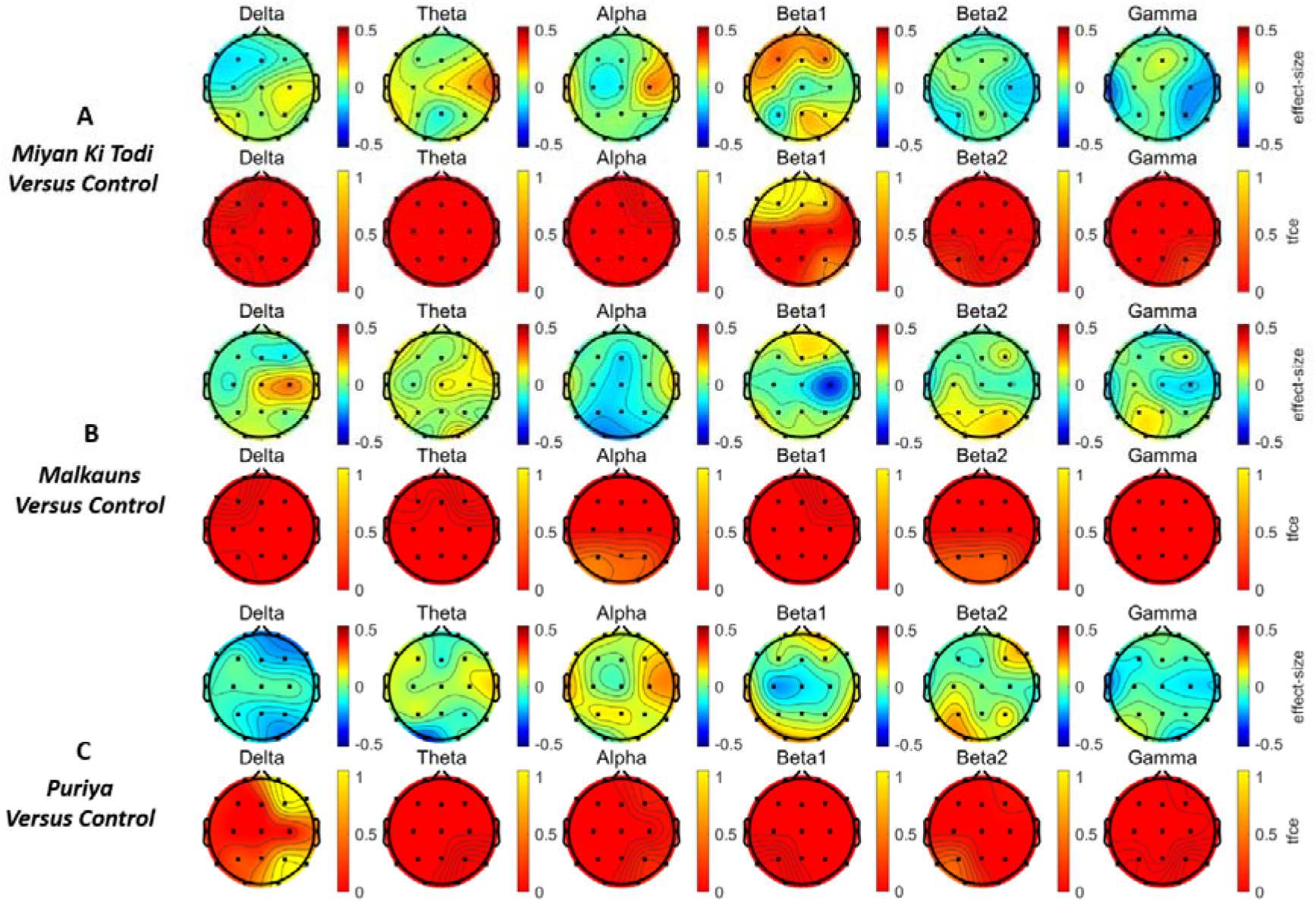
Scalp maps showing group differences in electrode-level band power between the intervention and control groups after intervention.

### Correlated component analysis (CorrCA) based on Inter Subject Correlation (ISC)

To explore the most temporally consistent EEG pattern for the different conditions and groups, we performed a Correlated Component Analysis (CorrCA), which extracts the components correlated between subjects. As our EEG segments were not well timed for intervention stimuli, we used frequency domain data (average power spectral data within each 10 min condition) for CorrCA. Based on the spectral distribution of the first three most correlated components, the first component is globally distributed low-frequency activity (C1), the second component represents posterior dominant alpha-beta1 activity (C2), and the third component represents peripherally dominant broad-band activity (C3) (Fig 7).

**Figure 7:**
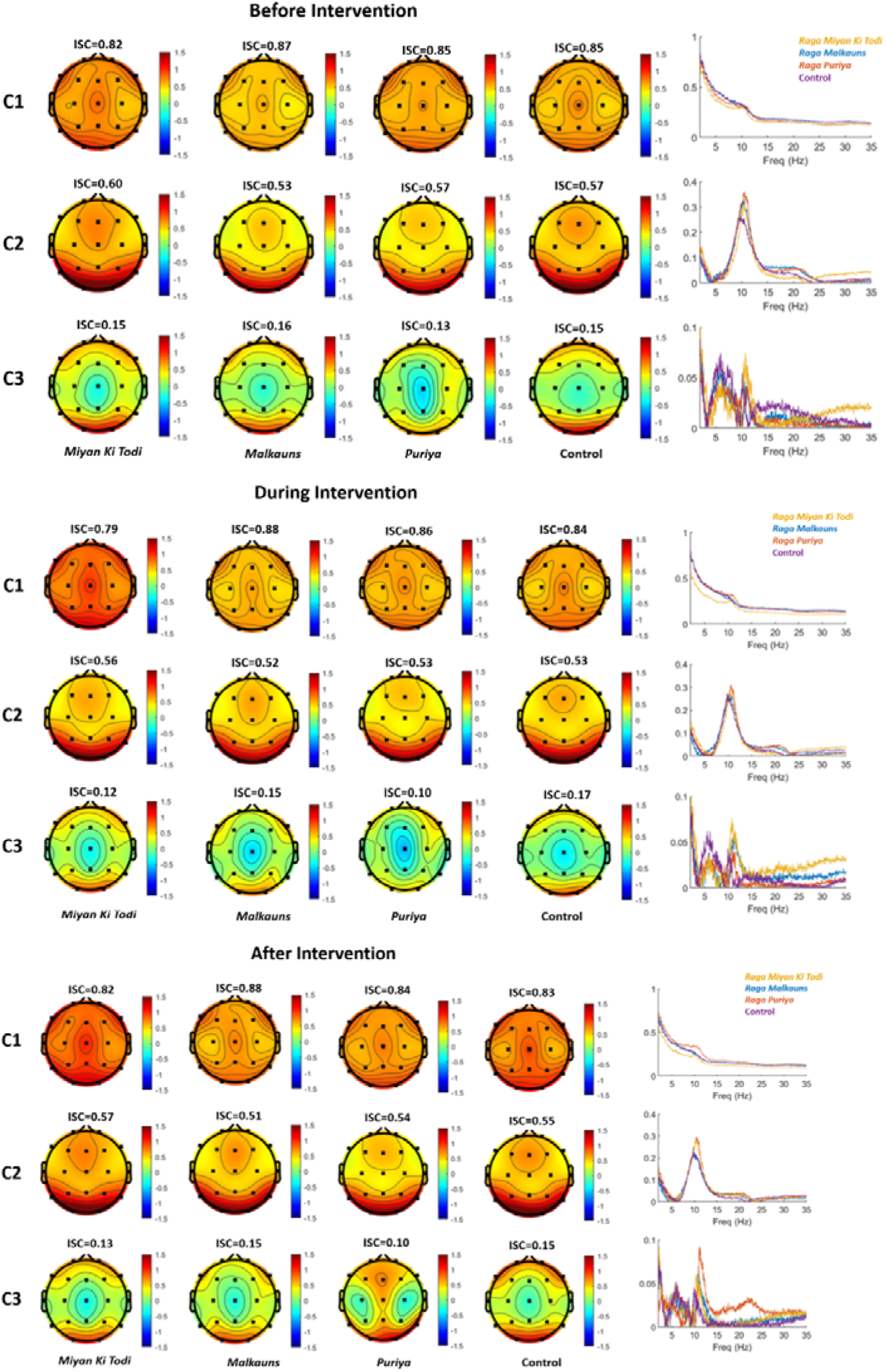
Scalp distribution and spectral pattern of the first three components (C1, C2 & C3) based on CorrCA before (7a), during (7b), and after (7c) intervention.

In terms of the spectral dynamics of the CorrCA components, both *raga Malkauns* and *raga Miyan ki Todi* groups showed a similar pattern of decrease in C1 power and increase in C2 power during intervention relative to baseline when compared to the control group (Fig 8). Even after the intervention, this pattern was strong for *raga Malkauns*, but weaker for *raga Miyan ki Todi*. Whereas *raga Puriya* showed only a weak decrease in C1 (after intervention), compared to the control group.

**Figure 8:**
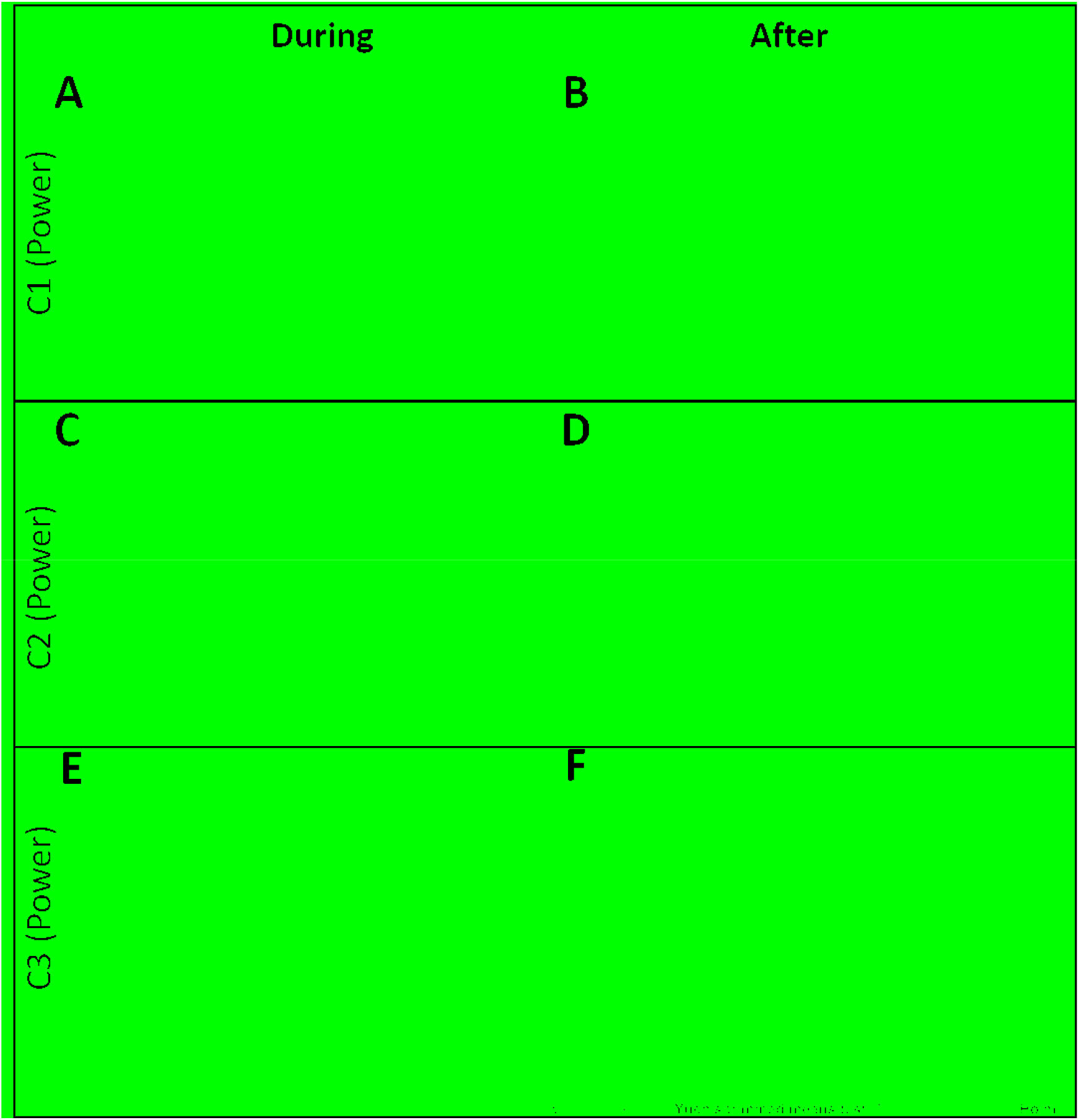
Change in the total spectral power of the components (C1, C2, & C3) between the four groups during and after the intervention, relative to before the intervention [C1 during (8A) C1 after (8B) C2 during (8C) C2 after (8D) C3 during (8E) C3 after (8F) intervention].

ISC scores were comparable between groups, except for *raga Puriya, which showed* a marginal drop in C3 after intervention (Fig 9).

**Figure 9:**
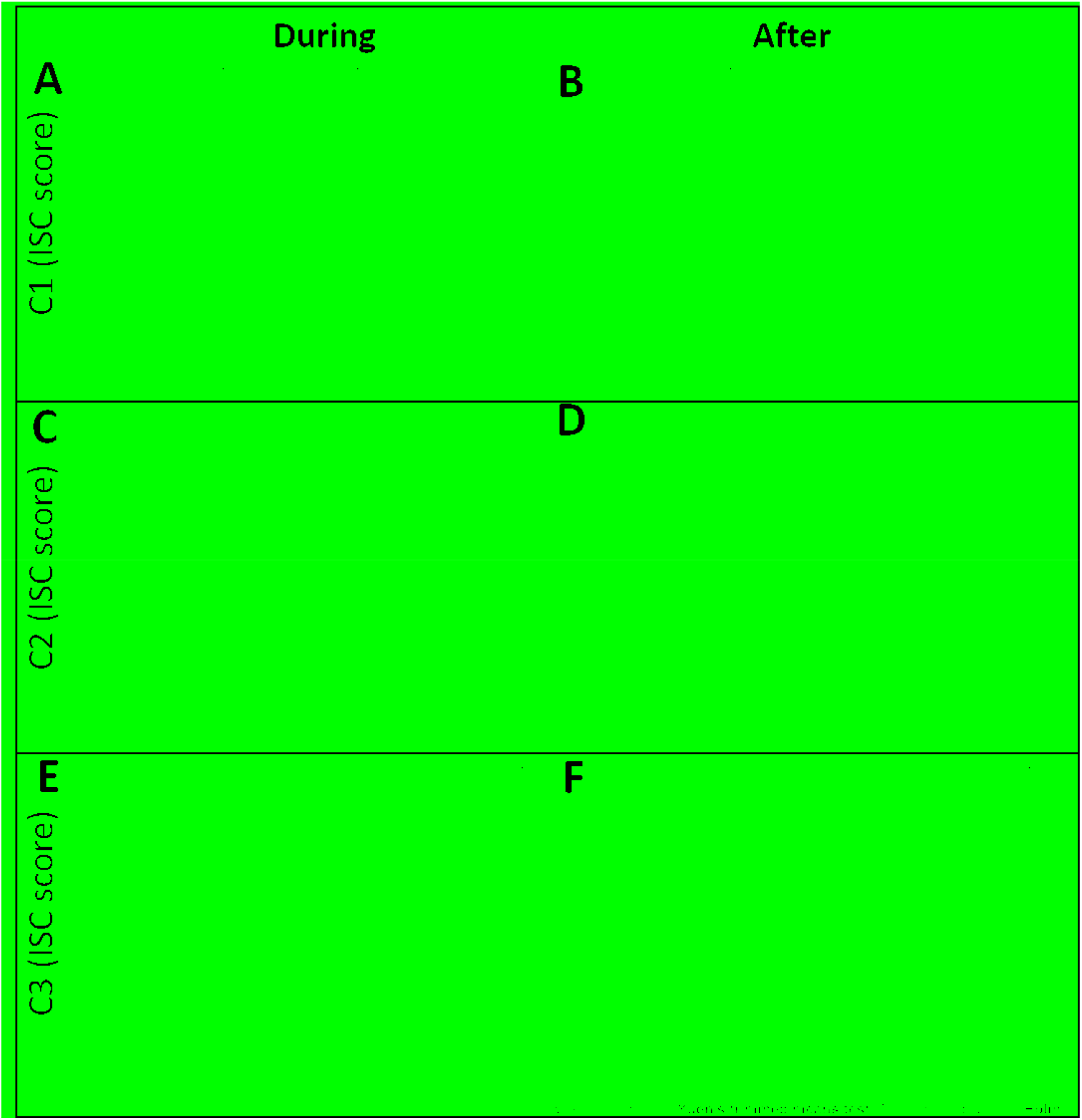
Change in the inter-subject correlation scores of the components (C1, C2, & C3) between the four groups during and after the intervention, relative to before the intervention [C1 during (8A) C1 after (8B) C2 during (8C) C2 after (8D) C3 during (8E) C3 after (8F) intervention].

## Discussion

In this study, we evaluated the spectral effect of EEG of acoustic stimuli (three Indian music and one control stimulus) in healthy young participants, the intervention lasting 10 minutes. The three musical stimuli consisted of instrumental music based on different *ragas* of Indian classical music (*raga Miyan ki Todi, raga Malkauns, and raga Puriya*) that were chosen based on Indian music ancient literature (52,53). The sociodemographic data, anxiety, and autonomic changes pre, during and post-intervention has been explained in (34). In the electrode-level analysis of EEG, during the intervention, the *raga Malkauns* group showed a significant increase in left frontal gamma power. After the intervention, the *Puriya raga* group showed a decrease in the right frontoparietal delta power, and the *Miyan ki Todi raga* group showed a frontal increase in beta1 power. Exploring this further, the component-level analysis showed a decrease in C1 power (globally distributed low-frequency activity) and an increase in C2 power (posterior dominant alpha-beta1 activity) with *raga Malkauns* (strong both during and after intervention) and *raga Miyan ki Todi* (strong during and weak after intervention), whereas *raga Puriya* showed only a weak decrease in C1 (after intervention), compared to control group.

### EEG power spectral patterns

On electrode-level power spectral analysis, we saw a decrease in delta power and an increase in beta1 and gamma power, and no significant change in alpha power, in music intervention groups relative to the control group.

Several EEG studies report conflicting evidence of electrode-level power spectral changes (decrease or increase, or null responses) when listening to music. For example, an EEG study that used Mozart’s K.448 music (54), reported a significant drop in alpha power during listening globally (F3-C3, F4-C4, C3-T3, C4-T4, T3-O1, and T4-O2) which persisted in posterior sites (T3-O1, T4-O2, O1-C3, and O2-C4) post-music, compared to pre-music alpha power. This study also observed a significant decrease in theta (at T3-O1 & O1-C3) and beta (at T3-O1, O1-C3 & O2-C4) power during listening, which partially persisted after music. Alpha power is often associated with active inhibition and therefore a drop in alpha power could indicate an overall increase in the level of brain activation (or disinhibition) that occurred when listening to the acoustic stimulus (55) or when actively participating in processing or anticipating a stimulus (56). But since alpha power drop co-occurred with drop in closer spectral bands (theta and beta) while listening to music, the spectral specificity mentioned before may not be true. Another EEG study that involved listening to popular classical symphonic pieces has shown that the alpha power increased in the parietal and occipital areas of both hemispheres during listening and that the maximum frequency of the alpha band was significantly reduced (57). The authors concluded that the level of brain activation was reduced on listening to music.

A previous study observed a reduction in delta power after slow-wave sleep brain-wave music, and this the authors interpreted to be a positive effect on sleep quality (58). Alternatively, conflicting evidence exists that this reduced delta could be due to lesser sleep (59). Listening to natural music has been found to drive the beta, theta, and delta activity (60), especially when the rhythm of the music falls in these frequency ranges. A MEG study of responses to pain induced during listening to preferred music versus personalized entrainment music found that preferred music reduced delta power in the cingulate gyrus, while entrainment music led to changes in gamma power in the somatosensory regions (61). The drop in delta power after stopping *raga Puriya* in the current study could be due to alertness or divergent thinking after stopping the intervention (62). Other frequency bands found to change when listening to a Mozart musical piece, Sonata K.448, were reduced theta power in the left temporal area; increased beta in the left and right temporal, the left frontal, and increased alpha1 power in the left temporal region (63). It was found that the power of low-frequency brain waves increased in the auditory cortex with a gradual increase in theta and alpha power in the amygdala and orbitofrontal cortex (probable higher analysis of music) with time along with an increase in the power of alpha, theta and beta1 waves in the orbitofrontal cortex while listening to consonant sounds (31). Taken together, there could be a globally distributed alpha activity that decreases along with other frequency bands (like delta, theta, and beta) and a more posterior dominant alpha activity that increases while listening to engaging music. This is possibly what our component-level analysis captured as C1 power decreases and C2 power increases during and after music listening. Similarly, studies have found a co-occurrence of power changes in higher frequency bands. In one study, enhanced spatial performance after exposure to music was associated with a lower alpha 2 (10.5-11.97 Hz) and a higher beta 1 (12.02-17.97 Hz) (63). In another study, there was a uniform reduction of alpha power and an increase in the gamma high power localized in the electrodes over or nearby the auditory-cortex brain regions (e.g. FT7, TP7, FC3, FT8, T4), during music listening (47). This may be captured by the slight increase in C3 power during listening in the *raga Malkauns* group.

Alpha activity in EEG often increases with an increase in task demands (for example, answering questions about stimuli) (64). The beta and alpha rhythms are seen in the awake state and in that, beta rhythm is usually associated with increased alertness, cortical integrity, stress, strong emotions, and cognitive processes (65,66). As reported before, a significant increase in EEG beta power during music listening, positively correlated with regional cerebral blood flow (17). This was proposed to be due to active cognitive sound processing, within the premotor-posterior parietal framework. Similar to beta, an increase in gamma activity is associated with selective attention, working memory, and conscious stimulus recognition (67,68). Beta rhythms are shown to predict listener-specific neural signature in naturalistic music listening (69). Gamma activity has been implicated in the perceptual binding of musical features at the sensory level and the matching of external acoustic information to internal thought processes to form meaningful concepts (70,71). Gamma is also found to be higher in trained musicians, reflecting improved binding of musical features (72–74) and may be related to musical expectations (75). A lower gamma event-related synchronization was recorded among dancers, during listening to preferred music, said to be due to selective attention to stimuli while probably planning/imagining dance movements. Alternatively, the rise in gamma and beta was postulated to be due to familiar music, which increases emotional provocation. (76). The same study also showed a significant decrease in alpha activity for their expertise-related music compared to other music, probably due to inhibition release (77) and cortical involvement in listening to their own music. Therefore, the decrease in globally distributed low-frequency activity (C1), the increase in posterior dominant alpha-beta1 activity (C2), and the weak increase in peripherally dominant broad-band activity that includes beta/gamma (C3), shown most prominently by the group *raga Malkauns, (*and insignificantly by *raga Miyan ki Todi)* could be due to attention modulation, increased alertness, binding of music features or may also serve as a reliable indicator of liking to music (78).

### Significance of *Component-Level Analysis Based on Intersubject Correlation*

In the current study, ISC scores were comparable between the groups, except with *raga Puriya* which showed a marginal drop in C3 power (peripherally dominant broad-band activity) after the intervention. It is important to note that all the participants were Indians, and the music was exclusively recorded for the current study (not familiar with the clippings used). But, we cannot exclude the participants’ prior exposure to these tones or combinations of tones. It was previously shown that ISC is capable of tracking musical engagement even though the behavior is not recorded. Our hypothesis that repetition of the phrases may reduce ISC was partially true only for the group that listened to *raga Puriya*.

ISC examines shared brain responses between participants, the degree to which their responses match each other, and musical stimuli, it represents the degree to which the music is gripping their brains and is driving their experience (30). The ISC of EEG has been used as an index of engagement with naturalistic stimuli such as movies, stories, speeches, and music. The ISC has both lower-level processes, such as sensory processing, and higher-level processes, such as memory retention (24,26). However, ISC is shown to be affected by attention (25,79). When attention is diverted away from sensory processes, the resulting ISC is shown to be smaller among participants (79,80). Thus, when attention is allocated to the stimulus, ISC is increased. In a previous study, the ISC of the relative component (RC1) in EEG was highest for a foreign language narrative and lowest for an English narrative among fluent English speakers (27). ISC is shown to be higher with familiar music, compared to unfamiliar styles. Furthermore, the repetition of familiar music leads to a decrease in ISC indicating a reduction in engagement (30).

As our EEG segments were not very well time-locked to intervention stimuli, we used frequency domain data (average power spectral data within each 10 min condition) for CorrCA, based on the spectral distribution of the first three most correlated components. In the current study, we used the first three components with the highest ISC values in each group, and their ISC values were statistically comparable and showed similar patterns. This means that we captured spatiospectral components that showed comparable engagement during the session and their spectral pattern might better represent the intervention-related changes. Previous studies have observed that passive listening to natural music evoked significant intersubject and stimulus-response correlations, suggesting distinct neural correlates of musical engagement (81). Another recent study also showed significant ISC during periods of escalated tension with natural music (cello, strings) but at the high point, there was no significance (82). These authors have also recorded that EEG ISCs are highest for the remix stimulus that had more attention-catching musical events and lowest for our most repetitive manipulation, Tremolo (83).

Our EEG spectral findings, especially those from component-level analysis, align with findings related to default mode network (DMN) activity and its relation to imagery or self-referential thoughts. To support this notion, a high-density EEG study that intermittently switched the attention of participants from internal (autobiographical remembering) to external processing (GO-NO-GO task) processing, found that autobiographical remembering was associated with an increase in spectral power in alpha and beta and a decreased in the delta band (84). At the source level, the alpha power increase was localized to regions of DMN. The decrease in delta power is more pronounced when the autobiographical contents have positive emotions. Furthermore, the more posteriorly distributed alpha increase would suggest a relatively higher activity in the anterior DMN hub, which is involved in mostly conscious modeling, planning, and control functions, and relatively lower activity in the posterior hub, which is involved in mostly unconscious processes that include self-representation, emotion, and salience detection (85). However, there are some methodological limitations in this study. Due to the lower density of the EEG electrodes, we cannot verify the relation of our finding to the DMN activity through source localization analysis. Furthermore, we had not collected phenomenological reports after the intervention sessions that could be subjected to structured analysis and correlated with the EEG findings. These limitations should be addressed in subsequent studies.

To corroborate these findings with autonomic changes and anxiety levels, all three music interventions were found to reduce state anxiety levels, along with a reduction in salivary alpha-amylase (34). During the intervention, the *raga Miyan ki Todi and raga Puriya* groups had a significant drop in the parasympathetic parameters of heart rate variability (HRV), while after the intervention these two modes led to a significant increase in the parasympathetic response. In contrast, the *raga Malkauns* group showed a sustained rise in parasympathetic tone, similar to that seen in the control group. However, *raga Malkauns* caused a significant reduction in anxiety levels that was not observed in the control group. Though we did not find any difference in the visual analog scale for the liking of the music, it may be observed that functional neural plasticity could occur even when not subjectively perceived. In this study, the electrocardiogram data, from which HRV was derived was computed for a minimum of 5 minutes to a maximum of 10 minutes for each condition, which meant that the participants had heard most of the phrases of the musical piece (EEG was computed for 3 minutes duration in the current manuscript). Furthermore, it would be interesting to understand whether these changes were modified after controlling for baseline data from the control group, as seen in the current EEG study.

To our knowledge, this is the first time that three acoustic stimuli in the form of Indian melodic modes have been studied systematically and scientifically as acoustic interventions for their short-term neuroplastic effect on the EEG power spectrum, with ISC-based component-level analysis, among a comparatively larger sample of healthy young individuals. Reduction in globally distributed low-frequency activity and increase in posterior dominant alpha-beta1 activity may be characteristic of passive listening to relaxing Indian modes, which may persist even after 10 minutes of the listening period. Among the modes, *raga Malkauns* showed this effect most prominently, followed by *raga Miyan ki Todi* and least by *raga Puriya*. As the increase in posterior alpha and low beta power is associated with DMN activity and a decrease in delta power with positive emotional memory, the spectral pattern we observed may indicate the observation of positive autobiographical memory while listening to musical modes and thus contribute to a relaxing experience. Further studies may include phenomenological reports to support these findings, and build a stronger scientific foundation for the use of music in medicine. As ISC-based brain activity is modulated by training, studies may try to explore the effect of musical training and genre familiarity (30) aspects. Different musical stimuli that are known to be emotionally stimulating can be studied, as ISC is said to vary with time-based emotional stimuli such as stories or movies. To exactly know the neural substrates activated within and between participants passively listening to the different scales, it is better to use higher-density EEG or fMRI data.

## Author Note

The first author was previously affiliated with Ramaiah Medical College, MSR Nagar, MSRIT Post, Bangalore 560054, Karnataka, India, where the data collection was done.

## Author Abbreviations

Kirthana Kunikullaya U-KKU, Arun Sasidharan-SA, Vijayadas-V, Radhika Kunnavil-KR, Jaisri Goturu-JG, Nandagudi Srinivasa Murthy-MSN

## Author Contributions

Conceptualization, Funding acquisition—KKU; Data curation— KKU; Formal analysis—SA, KKU; Investigation—KKU, V; Methodology—KKU, V, KR, JG, MSN; Project administration—KKU, JG, MSN; Resources—KKU, V, KR; Software— KKU, SA; Supervision—JG, MSN; Validation— KKU, SA, KR; Writing original draft— KKU, SA; Review and editing—KKU, SA, V, KR, JG, MSN. All authors have approved the final version of the manuscript and agree to be accountable for all aspects of the work. All persons designated as authors qualify for authorship, and all those who qualify for authorship are listed.

## Data Availability Statement

The data that support the findings of this study are available on request from the corresponding author. The data are not publicly available due to privacy or ethical restrictions.

## Acknowledgments

The authors acknowledge funding agencies and the exclusive recording shared by Vidhwan Pandit Pravin Godkhindi, an eminent flutist, for this study. We thank Mamta S Vernekar, J Sundaramma, and Anjani Bhushan for their assistance with data collection. We thank Chaitra L and Sindhu Reddy for their assistance with time marking. We like to thank all the volunteers who participated in the study.

## References

1. Kaplan S. The restorative benefits of nature: Toward an integrative framework. Journal of Environmental Psychology. 1995 Sep 1;15(3):169–82.

2. Seymour V. The Human–Nature Relationship and Its Impact on Health: A Critical Review. Frontiers in Public Health [Internet]. 2016 [cited 2022 Mar 31];4. Available from: https://www.frontiersin.org/article/10.3389/fpubh.2016.00260

3. Schreuder E, van Erp J, Toet A, Kallen VL. Emotional Responses to Multisensory Environmental Stimuli: A Conceptual Framework and Literature Review. SAGE Open. 2016 Jan 1;6(1):2158244016630591.

4. Diaz FM. Listening and Musical Engagement: An Exploration of the Effects of Different Listening Strategies on Attention, Emotion, and Peak Affective Experiences. Update: Applications of Research in Music Education. 2015 May 1;33(2):27–33.

5. Gustavson DE, Coleman PL, Iversen JR, Maes HH, Gordon RL, Lense MD. Mental health and music engagement: review, framework, and guidelines for future studies. Transl Psychiatry. 2021 Jun 22;11(1):1–13.

6. Schellenberg EG. Music and Cognitive Abilities. Curr Dir Psychol Sci. 2005 Dec 1;14(6):317–20.

7. Menon V, Levitin DJ. The rewards of music listening: response and physiological connectivity of the mesolimbic system. Neuroimage. 2005 Oct 15;28(1):175–84.

8. Cowen AS, Fang X, Sauter D, Keltner D. What music makes us feel: At least 13 dimensions organize subjective experiences associated with music across different cultures. PNAS. 2020 Jan 28;117(4):1924–34.

9. Eerola T, Vuoskoski JK, Peltola HR, Putkinen V, Schäfer K. An integrative review of the enjoyment of sadness associated with music. Physics of Life Reviews. 2018 Aug 1;25:100–21.

10. Kunikullaya KU, Goturu J, Muradi V, Hukkeri PA, Kunnavil R, Doreswamy V, et al. Combination of music with lifestyle modification versus lifestyle modification alone on blood pressure reduction - A randomized controlled trial. Complement Ther Clin Pract. 2016 May;23:102–9.

11. Kunikullaya Ubrangala K, Kunnavil R, Goturu J, M V, Prakash VS, Murthy NS. Effect of specific melodic scales of Indian music in reducing state and trait anxiety: A randomized clinical trial. Psychology of Music. 2021 Dec 23;03057356211055509.

12. Panteleeva Y, Ceschi G, Glowinski D, Courvoisier DS, Grandjean D. Music for anxiety? Meta-analysis of anxiety reduction in non-clinical samples. Psychology of Music. 2018 Jul 1;46(4):473–87.

13. DROIT-VOLET S, Ramos danilo, Bueno L, Bigand E. Music, emotion, and time perception: the influence of subjective emotional valence and arousal? Frontiers in Psychology [Internet]. 2013 [cited 2022 Feb 25];4. Available from: https://www.frontiersin.org/article/10.3389/fpsyg.2013.00417

14. Kunikullaya KU, Goturu J, Muradi V, Hukkeri PA, Kunnavil R, Doreswamy V, et al. Music versus lifestyle on the autonomic nervous system of prehypertensives and hypertensives—a randomized control trial. Complementary Therapies in Medicine. 2015 Oct 1;23(5):733–40.

15. de Witte M, Spruit A, van Hooren S, Moonen X, Stams GJ. Effects of music interventions on stress-related outcomes: a systematic review and two meta-analyses. Health Psychology Review. 2020 Apr 2;14(2):294–324.

16. Kirthana Kunikullaya U, Sasidharan A, Srinivasa R, Goturu J, Murthy NS. Temporal changes in electroencephalographic power spectrum on passive listening to three selected melodic scales of Indian music on healthy young individuals - a randomized controlled trial. Music and Medicine. 2022 Jan 31;14(1):06–26.

17. Nakamura S, Sadato N, Oohashi T, Nishina E, Fuwamoto Y, Yonekura Y. Analysis of music-brain interaction with simultaneous measurement of regional cerebral blood flow and electroencephalogram beta rhythm in human subjects. Neurosci Lett. 1999 Nov 19;275(3):222–6.

18. Fletcher PC, Frith CD, Baker SC, Shallice T, Frackowiak RS, Dolan RJ. The mind’s eye--precuneus activation in memory-related imagery. Neuroimage. 1995 Sep;2(3):195–200.

19. Psychophysiological responsivity to Indian instrumental music - Uma Gupta, B. S. Gupta, 2005 [Internet]. [cited 2019 May 17]. Available from: https://journals.sagepub.com/doi/10.1177/0305735605056144

20. Maity AK, Pratihar R, Mitra A, Dey S, Agrawal V, Sanyal S, et al. Multifractal Detrended Fluctuation Analysis of alpha and theta EEG rhythms with musical stimuli. Chaos, Solitons & Fractals. 2015 Dec 1;81:52–67.

21. Lee EJ, Bhattacharya J, Sohn C, Verres R. Monochord sounds and progressive muscle relaxation reduce anxiety and improve relaxation during chemotherapy: a pilot EEG study. Complement Ther Med. 2012 Dec;20(6):409–16.

22. Gruskin DC, Patel GH. Brain connectivity at rest predicts individual differences in normative activity during movie watching. NeuroImage. 2022 Jun 1;253:119100.

23. Nastase SA, Gazzola V, Hasson U, Keysers C. Measuring shared responses across subjects using intersubject correlation. Social Cognitive and Affective Neuroscience. 2019 Aug 7;14(6):667–85.

24. Hasson U, Nir Y, Levy I, Fuhrmann G, Malach R. Intersubject Synchronization of Cortical Activity During Natural Vision. Science. 2004 Mar 12;303(5664):1634–40.

25. Dmochowski J, Sajda P, Dias J, Parra L. Correlated Components of Ongoing EEG Point to Emotionally Laden Attention – A Possible Marker of Engagement? Frontiers in Human Neuroscience [Internet]. 2012 [cited 2022 Mar 31];6. Available from: https://www.frontiersin.org/article/10.3389/fnhum.2012.00112

26. Cohen SS, Henin S, Parra LC. Engaging narratives evoke similar neural activity and lead to similar time perception. Sci Rep. 2017 Jul 4;7(1):4578.

27. Ki JJ, Kelly SP, Parra LC. Attention Strongly Modulates Reliability of Neural Responses to Naturalistic Narrative Stimuli. J Neurosci. 2016 Mar 9;36(10):3092–101.

28. Kauppi JP, Jääskeläinen I, Sams M, Tohka J. Inter-subject correlation of brain hemodynamic responses during watching a movie: localization in space and frequency. Frontiers in Neuroinformatics [Internet]. 2010 [cited 2022 Mar 31];4. Available from: https://www.frontiersin.org/article/10.3389/fninf.2010.00005

29. Kaneshiro B, Dmochowski JP, Norcia AM, Berger J. TOWARD AN OBJECTIVE MEASURE OF LISTENER ENGAGEMENT WITH NATURAL MUSIC USING INTER-SUBJECT EEG CORRELATION.:5.

30. Madsen J, Margulis EH, Simchy-Gross R, Parra LC. Music synchronizes brainwaves across listeners with strong effects of repetition, familiarity and training. Scientific Reports [Internet]. 2019 Dec [cited 2020 May 13];9(1). Available from: http://www.nature.com/articles/s41598-019-40254-w

31. Schaefer RS, Vlek RJ, Desain P. Music perception and imagery in EEG: alpha band effects of task and stimulus. Int J Psychophysiol. 2011 Dec;82(3):254–9.

32. Janata P, Birk JL, Van Horn JD, Leman M, Tillmann B, Bharucha JJ. The Cortical Topography of Tonal Structures Underlying Western Music. Science. 2002 Dec 13;298(5601):2167–70.

33. Bowling DL, Sundararajan J, Han S, Purves D. Expression of Emotion in Eastern and Western Music Mirrors Vocalization. PLoS One. 2012 Mar 14;7(3):e31942.

34. Kunikullaya Ubrangala K, Kunnavil R, Sanjeeva Vernekar M, Goturu J, Vijayadas, Prakash VS, et al. Effect of Indian Music as an Auditory Stimulus on Physiological Measures of Stress, Anxiety, Cardiovascular and Autonomic Responses in Humans—A Randomized Controlled Trial. European Journal of Investigation in Health, Psychology and Education. 2022 Oct;12(10):1535–58.

35. Graff V, Cai L, Badiola I, Elkassabany NM. Music versus midazolam during preoperative nerve block placements: a prospective randomized controlled study. Reg Anesth Pain Med. 2019 Aug 1;44(8):796–9.

36. Mathur A, Vijayakumar SH, Chakrabarti B, Singh NC. Emotional responses to Hindustani raga music: the role of musical structure. Front Psychol. 2015;6:513.

37. Watanabe K, Ooishi Y, Kashino M. Heart rate responses induced by acoustic tempo and its interaction with basal heart rate. Sci Rep. 2017 07;7:43856.

38. Liu Y, Liu G, Wei D, Li Q, Yuan G, Wu S, et al. Effects of Musical Tempo on Musicians’ and Non-musicians’ Emotional Experience When Listening to Music. Front Psychol. 2018;9:2118.

39. Levitin DJ, Grahn JA, London J. The Psychology of Music: Rhythm and Movement. Annu Rev Psychol. 2018 04;69:51–75.

40. Zhao TC, Lam HTG, Sohi H, Kuhl PK. Neural processing of musical meter in musicians and non-musicians. Neuropsychologia. 2017 Nov;106:289–97.

41. Reinhardt U. [Investigations into synchronisation of heart rate and musical rhythm in a relaxation therapy in patients with cancer pain]. Forsch Komplementarmed. 1999 Jun;6(3):135–41.

42. Stith CC. The Effects of Musical Tempo and Dynamic Range on Heart Rate Variability in Healthy Adults: A Counterbalanced, Within-subjects Study. 2015 May;100.

43. Patel AD, Iversen JR. The evolutionary neuroscience of musical beat perception: the Action Simulation for Auditory Prediction (ASAP) hypothesis. Front Syst Neurosci. 2014;8:57.

44. Daly I, Hallowell J, Hwang F, Kirke A, Malik A, Roesch E, et al. Changes in music tempo entrain movement related brain activity. Conf Proc IEEE Eng Med Biol Soc. 2014;2014:4595–8.

45. Schaub K, Demos L, Centeno T, Daugherty B. Effects of Musical Tempo on Heart Rate, Brain Activity, and Short-term Memory. Journal of Advanced Student Science (JASS). 2011;(1):11.

46. Idrobo-Ávila EH, Loaiza-Correa H, van Noorden L, Muñoz-Bolaños FG, Vargas-Cañas R. Different Types of Sounds and Their Relationship With the Electrocardiographic Signals and the Cardiovascular System - Review. Front Physiol. 2018;9:525.

47. Adamos DA, Laskaris NA, Micheloyannis S. Harnessing functional segregation across brain rhythms as a means to detect EEG oscillatory multiplexing during music listening. Journal of Neural Engineering. 2018 Jun 1;15(3):036012.

48. EEGLAB [Internet]. [cited 2020 Jan 5]. Available from: https://sccn.ucsd.edu/eeglab/index.php

49. Chang CY, Hsu SH, Pion-Tonachini L, Jung TP. Evaluation of Artifact Subspace Reconstruction for Automatic Artifact Components Removal in Multi-Channel EEG Recordings. IEEE Transactions on Biomedical Engineering. 2020 Apr;67(4):1114–21.

50. Mair P, Wilcox R. Robust statistical methods in R using the WRS2 package. Behav Res Methods. 2020;52(2):464–88.

51. Holm S. A Simple Sequentially Rejective Multiple Test Procedure. Scandinavian Journal of Statistics. 1979;6(2):65–70.

52. McNeil A. Ragas, Recipes, and Rasas [Internet]. Oxford Handbooks Online. 2015 [cited 2020 Apr 29]. Available from: https://www.oxfordhandbooks.com/view/10.1093/oxfordhb/9780199935321.001.0001/oxfordhb-9780199935321-e-43

53. Jairazbhoy NA. The Rāgs of North Indian Music: Their Structure and Evolution. Popular Prakashan; 1995. 252 p.

54. Lin LC, Ouyang CS, Chiang CT, Wu RC, Wu HC, Yang RC. Listening to Mozart K.448 decreases electroencephalography oscillatory power associated with an increase in sympathetic tone in adults: a post-intervention study. JRSM Open. 2014 Oct 8;5(10):2054270414551657.

55. Mikutta C, Altorfer A, Strik W, Koenig T. Emotions, Arousal, and Frontal Alpha Rhythm Asymmetry During Beethoven’s 5th Symphony. Brain Topogr. 2012 Oct 1;25(4):423–30.

56. Weisz N, Hartmann T, Müller N, Lorenz I, Obleser J. Alpha Rhythms in Audition: Cognitive and Clinical Perspectives. Front Psychol. 2011 Apr 26;2:73.

57. Sulimov AV, Liubimova IV, Pavlygina RA, Davydov VI. [Spectral analysis of the human EEG while listening to music]. Zh Vyssh Nerv Deiat Im I P Pavlova. 2000 Feb;50(1):62–7.

58. Gao D, Long S, Yang H, Cheng Y, Guo S, Yu Y, et al. SWS Brain-Wave Music May Improve the Quality of Sleep: An EEG Study. Front Neurosci. 2020 Feb 11;14:67.

59. Johnson JM, Durrant SJ. Commentary: SWS Brain-Wave Music May Improve the Quality of Sleep: An EEG Study. Front Neurosci. 2021 Feb 1;15:609169.

60. Tichko P, Kim JC, Large E, Loui P. Integrating music-based interventions with Gamma-frequency stimulation: Implications for healthy ageing. European Journal of Neuroscience. 2022;55(11–12):3303–23.

61. Hauck M, Metzner S, Rohlffs F, Lorenz J, Engel AK. The influence of music and music therapy on pain-induced neuronal oscillations measured by magnetencephalography. Pain. 2013 Apr;154(4):539–47.

62. Boot N, Baas M, Mühlfeld E, de Dreu CKW, van Gaal S. Widespread neural oscillations in the delta band dissociate rule convergence from rule divergence during creative idea generation. Neuropsychologia. 2017 Sep;104:8–17.

63. Rideout BE, Laubach CM. EEG correlates of enhanced spatial performance following exposure to music. Percept Mot Skills. 1996 Apr;82(2):427–32.

64. Cooper NR, Croft RJ, Dominey SJJ, Burgess AP, Gruzelier JH. Paradox lost? Exploring the role of alpha oscillations during externally vs. internally directed attention and the implications for idling and inhibition hypotheses. Int J Psychophysiol. 2003 Jan;47(1):65–74.

65. Sammler D, Grigutsch M, Fritz T, Koelsch S. Music and emotion: Electrophysiological correlates of the processing of pleasant and unpleasant music. Psychophysiology. 2007 Mar;44(2):293–304.

66. Kozelka JW, Pedley TA. Beta and mu rhythms. J Clin Neurophysiol. 1990 Apr;7(2):191–207.

67. Pavlova M, Birbaumer N, Sokolov A. Attentional modulation of cortical neuromagnetic gamma response to biological movement. Cereb Cortex. 2006 Mar;16(3):321–7.

68. Carozzo S, Garbarino S, Serra S, Sannita WG. Function-Related Gamma Oscillations and Conscious Perception. Journal of Psychophysiology. 2010 Jan;24(2):102–6.

69. Pandey P, Miyapuram KP, Lomas D. Non-Linear Features of β Brain Rhythms Predict Listener-Specific Neural Signature in Naturalistic Music Listening. 2022 Aug 22 [cited 2022 Nov 1]; Available from: https://www.techrxiv.org/articles/preprint/Non-Linear_Features_of_Brain_Rhythms_Predict_Listener-Specific_Neural_Signature_in_Naturalistic_Music_Listening/20502423/1

70. Bertrand O, Tallon-Baudry C. Oscillatory gamma activity in humans: a possible role for object representation. Int J Psychophysiol. 2000 Dec 1;38(3):211–23.

71. Rodriguez E, George N, Lachaux JP, Martinerie J, Renault B, Varela FJ. Perception’s shadow: long-distance synchronization of human brain activity. Nature. 1999 Feb 4;397(6718):430–3.

72. Bhattacharya J, Petsche H, Pereda E. Interdependencies in the spontaneous EEG while listening to music. Int J Psychophysiol. 2001 Nov;42(3):287–301.

73. Shahin AJ, Roberts LE, Chau W, Trainor LJ, Miller LM. Music training leads to the development of timbre-specific gamma band activity. Neuroimage. 2008 May 15;41(1):113–22.

74. Pallesen KJ, Bailey CJ, Brattico E, Gjedde A, Palva JM, Palva S. Experience Drives Synchronization: The phase and Amplitude Dynamics of Neural Oscillations to Musical Chords Are Differentially Modulated by Musical Expertise. PLOS ONE. 2015 Aug 20;10(8):e0134211.

75. Snyder JS, Large EW. Gamma-band activity reflects the metric structure of rhythmic tone sequences. Brain Res Cogn Brain Res. 2005 Jun;24(1):117–26.

76. Nakano H, Rosario MAM, de Dios C. Experience Affects EEG Event-Related Synchronization in Dancers and Non-dancers While Listening to Preferred Music. Frontiers in Psychology [Internet]. 2021 [cited 2022 Sep 8];12. Available from: https://www.frontiersin.org/articles/10.3389/fpsyg.2021.611355

77. Klimesch W. α-band oscillations, attention, and controlled access to stored information. Trends Cogn Sci. 2012 Dec;16(12):606–17.

78. Hadjidimitriou SK, Hadjileontiadis LJ. Toward an EEG-based recognition of music liking using time-frequency analysis. IEEE Trans Biomed Eng. 2012 Dec;59(12):3498–510.

79. Rosenkranz M, Holtze B, Jaeger M, Debener S. EEG-Based Intersubject Correlations Reflect Selective Attention in a Competing Speaker Scenario. Frontiers in Neuroscience [Internet]. 2021 [cited 2022 Apr 22];15. Available from: https://www.frontiersin.org/article/10.3389/fnins.2021.685774

80. Cohen SS, Madsen J, Touchan G, Robles D, Lima SFA, Henin S, et al. Neural engagement with online educational videos predicts learning performance for individual students. Neurobiology of Learning and Memory. 2018 Nov 1;155:60–4.

81. Kaneshiro B, Nguyen DT, Norcia AM, Dmochowski JP, Berger J. Natural music evokes correlated EEG responses reflecting temporal structure and beat. Neuroimage. 2020 Jul 1;214:116559.

82. Kaneshiro B, Nguyen DT, Norcia AM, Dmochowski JP, Berger J. Inter-Subject EEG Correlation Reflects Time-Varying Engagement with Natural Music [Internet]. bioRxiv; 2021 [cited 2022 Apr 22]. p. 2021.04.14.439913. Available from: https://www.biorxiv.org/content/10.1101/2021.04.14.439913v2

83. Dauer T, Nguyen DT, Gang N, Dmochowski JP, Berger J, Kaneshiro B. Inter-subject Correlation While Listening to Minimalist Music: A Study of Electrophysiological and Behavioral Responses to Steve Reich’s Piano Phase. Frontiers in Neuroscience [Internet]. 2021 [cited 2022 Apr 22];15. Available from: https://www.frontiersin.org/article/10.3389/fnins.2021.702067

84. Knyazev GG, Savostyanov AN, Bocharov AV, Dorosheva EA, Tamozhnikov SS, Saprigyn AE. Oscillatory correlates of autobiographical memory. International Journal of Psychophysiology. 2015 Mar;95(3):322–32.

85. Knyazev GG. Extraversion and anterior vs. posterior DMN activity during self-referential thoughts. Front Hum Neurosci [Internet]. 2013 [cited 2022 Oct 20];6. Available from: http://journal.frontiersin.org/article/10.3389/fnhum.2012.00348/abstract

