## Supplementary material for "Electroencephalographic power spectrum and intersubject correlation on acoustic stimulation with modes of Indian music: a randomized controlled trial": Supplemntary file

S1. Music intervention – Melodic scales in detail

***Miyan ki Todi*, the *raga* A of this study,** is a Hindustani classical raga that gave its name to the *Todi thaat*, one of the ten modes of Hindustani classical music, also known as *Darbari Todi*, and sometimes Shuddha Todi, is amongst the more popular morning ragas of Hindustani music. The scale of *Miyan ki Todi* is *Arohana:* S r g m d N S' or 'd 'N S r g m d N S' or S r g m d P, m d N S' or S r g m P, m d N S' and *Avarohana:* S' N d P m g r S or S' N d P m d m g r g r S. *Vadi* and *Samavadi* are *Komal Dha* and *Komal Ga*. *Re, ga,* and *dha* are intoned slightly low, and *tivra ma* is very sharp. Bhatkhande pronounces *Komal Dha* as *Vadi* (primarily dominant), but some musicians accord this status to *Komal Ga*. According to him, *Komal Ga and Komal Re*, are candidates for the status of *samvadi* (secondary dominant). *Todi* is a *Raga* of the late morning. The prescribed time for the raga is the first three-hour slot after sunrise. The equivalent *raga* in Carnatic music is *Shubhapantuvarali.* *Miyan ki Todi* predominantly is mostly pervaded by a pensive, mournful mood which is then relieved in the *drut* (faster tempo) part, by a festive piece, possibly to alleviate the heavy pathos in the earlier stages of rendering, though not always. The composition is such as to afford an artist of high caliber to mold it in either the inherent pensive mood or to entirely present a festive mood. Despite this, the raga has attained a decent presence in the classicist as well as romanticist genres of Hindustani music. The common phrases used in this scale are: N. N. S r g / r g r / r g M^ P or r g M^ g P/ g M^ d P / M^ g M^ d / N d P / d d  N S’ [or] M^ d N S’/ N S’ r’ g’ r’/ d N S’ r’ g’ / r’ g’ r’ S’/ N r’ N d P / M^ P d M^ g [or] N d M^ g / r g r S (1). Popular songs based on this *raga* are: *Bhini bhini bhor (Asha Bhosle's Album Dil Padosi Hai), aeri mai to prem diwani mera dard na jane koi (A meerabai bhajan from the movie – Meera), Watan pe jo fida hoga (movie – Phool bane angaare)*(2), *oora serabahude neenu (title track of T N Seetharam Kannada serial ‘magalu jaanaki’).*

***Raga Malkauns*** belongs to *Kalyan thaat*, and is a majestic and somewhat introverted pentatonic raga. Ma is the pivotal tone of this *raga* and the tone in which the first string of the *tanpura* is usually tuned. *Ga, Dha,* and *Ni* may slightly oscillate. *Malkauns* should be performed in a slow and dignified manner, and to bring out its ethos the notes should be linked by glides, in particular N / D, D / M, and M / G (3). Time: Late night, 12 - 3. *Aarohan* (ascending scale): S G M D N M D S*; *Avaraohan* (descending scale): S* N D M G M G S, D S; * indicates a higher (third) octave. The *Rishabh* and the *pancham* are skipped in the scale. It is an *audhav - audhav* (5 notes in ascent and descent of the scale) *vakra (nishad* is rarely employed in *avaroh)*. The *vaadi samavaadi swaras* for this raga are d and g. The *vishranti sthaan* for this scale are G; D; S'; - D; G;. Example of *sanchar* (move/ phrases / flow) through this *raga*, S; G M D G M G ; M G ; G S ; ,D ,D S ; ,N ,M ,D S ; S G M D ; G M G ; M D S' ; N M D ; G M M G ; S ; ,D ,D S;. It is this preponderance of the *tivra madhyam,* thus intense training is required to perform this raga. Time for best effects is between (12 night - 3 am): 3^rd^ *prahar* of the night (*Ragas* are divided into *prahaars* whereby each *raga* has a specific period of the day when it is performed). The popular Hindi film songs based on *raga Malkauns* include *Aaye Sur Ke Panchhi Aaye (Movie - Sur Sangam), Adha Hai Chandrama Raat Adhi and Tu Chhupi Hai Kahan (Navrang), Man Tarapat Hari Darshan Ko (Baiju Bawra)* (3)*.* **Malkauns was the *Raga* B in this study.**

***Raga Puriya*** is a major hexatonic raga (*Shadhav – Shadhav*) of Hindustani classical music, belonging to the *marwa thaat*. Best performed just after sunset (2^nd^ prahar of the night). What is common among all types of *Puriya raag* are *komal* (flat) *Re*, *shuddha* (natural) *Ga*, *tivra* (sharp) *Ma*, and *shuddha* (natural) *Ni.* Aarohan: N r G M D N r S and avarohan: S N D M G r S N or r N D M Gg, M G r S. *Pancham Varjya. Rishabh Komal, Madhyam Teevra*. Rest all *Shuddha Swaras*. Mandra Saptak Nishad is the Nyas swar in Puriya. Illustrative combinations are: N r G ; G r ,N ,D ,N; ,N ,M ,D S; G M D N; N M G; G M D G M G; r S; G M D N D S'; N r' N M G ; G M D G M G r S (4). In this *raga*, N-M and D-G *sangati* is observed. Nishad is often skipped in Aaroh like G M D N D S'. *Raag Puriya* is often referred to as king of night *ragas*. The rasa / emotions related to this raga are *Shanti* (equanimity/peace) and *Gambhir* (seriousness) (5). ***Puriya* was the *Raga* C in this study.** Pure *Puriya* has not been very commonly used for film music.

**References**

1.	Raja D. Deepak Raja’s world of Hindustani Music: Raga Miya-ki Todi.... reluctant differentiation [Internet]. Deepak Raja’s world of Hindustani Music. 2011 [cited 2020 Mar 19]. Available from: http://swaratala.blogspot.com/2011/04/raga-miya-ki-todi-reluctant.html

2.	Film Songs in Rag Mian Ki Todi [Internet]. [cited 2020 Mar 19]. Available from: https://chandrakantha.com/raga_raag/film_song_raga/mian_ki_todi.shtml

3.	The Raga Guide - Malkauns [Internet]. 2009 [cited 2020 Mar 19]. Available from: https://web.archive.org/web/20090620030217/http://www.wyastone.co.uk/nrl/world/raga/malkauns.html

4.	Raag Puriya - Indian Classical Music - Tanarang.com [Internet]. [cited 2020 Mar 19]. Available from: http://www.tanarang.com/english/puriya_eng.htm

5.	Puriya. In: Wikipedia [Internet]. 2019 [cited 2020 Mar 19]. Available from: https://en.wikipedia.org/w/index.php?title=Puriya&oldid=927072231
